# SimMapNet: A Bayesian Framework for Gene Regulatory Network Inference Using Gene Ontology Similarities as External Hint

**DOI:** 10.1101/2025.04.09.647936

**Authors:** Maryam Shahdoust, Rosa Aghdam, Mehdi Sadeghi

**Author notes:** these authors contributed equally (Maryam Shahdoust, Rosa Aghdam).

## Abstract

**Motivation:** Gene regulatory network (GRN) reconstruction is a fundamental challenge in computational biology, and is crucial for understanding gene interactions. In this study, we aim to incorporate Gene Ontology (GO) similarities into the construction of GRNs. Our key assumption is that genes with higher similarity in Molecular Function, Biological Process, or Cellular Component categories are more likely to be functionally related and, therefore, more likely to be connected in the network. We introduce `SimMapNet`, a Bayesian framework that estimates the precision matrix, which serves as the adjacency matrix in a Gaussian graphical model (GGM) for GRN inference. `SimMapNet` enhances network inference by integrating GO similarities, which inform the hyperparameters of the prior distribution through a kernel function, incorporating biological prior knowledge in a principled manner.

**Results:** We evaluate `SimMapNet` on three datasets: two datasets from the SOS DNA-repair response pathway in *Escherichia coli* and one dataset from *Drosophila melanogaster*. The results demonstrate the algorithm’s superior performance compared to state-of-the-art methods such as GLASSO, GENIE3, and KBOOST in terms of F1-score. `SimMapNet` has low time complexity, making it suitable for constructing large networks. Our simulation results confirm that `SimMapNet` is particularly well-suited for scenarios with limited sample sizes, where traditional methods often struggle.

**Availability and implementation:** The datasets and R package of `SimMapNet` are available in the github repository, https://github.com/maryam-shahdoust/SimMapNet.

## Introduction

Developing an integrative framework for Gene Regulatory Network (GRN) that combines multiple data sources, machine learning algorithms, and network-based computational strategies remains a crucial challenge in the field, requiring continuous innovation and interdisciplinary collaboration [1]. Decades of biomedical research have led to the accumulation of extensive biological knowledge, including pathway information, transcription factor-target interactions, gene ontology (GO) annotations and other functional information, which are now available in public databases [2, 3]. Leveraging prior biological data can significantly enhance the statistical power and interpretability of GRN reconstruction, particularly in relation to complex phenotypes. However, traditional network inference methods rely solely on gene expression data [4, 5], often resulting in high false positive and false negative rates in the inferred network due to noise and limited coverage [6]. Studies have demonstrated that incorporating prior biological knowledge can refine network structure, reduce uncertainty, and improve predictive power [7, 8, 9, 10]. For instance, the Prior Lasso (pLasso) method partitions gene interactions based on pathway knowledge [10] and PriorPC that uses soft priors that assign to edges a probability of existence [8]. More recent approaches, such as GRNPT, integrate transformer-based embeddings from large-scale biological data to capture regulatory patterns from single-cell RNA sequencing [11].

In this study, we incorporate GO similarities into the construction of GRNs. Our key assumption is that genes with higher similarity in the Molecular Function (MF), Biological Process (BP) categories are more likely to be functionally related and, therefore, more likely to be connected in the network. This assumption is biologically motivated, as genes involved in similar biological processes or sharing molecular functions often participate in the same regulatory pathways [12, 13, 14]. Additionally, we also utilize Cellular Component (CC) similarities to assess the impact of different types of external information in improving the accuracy and robustness of GRN construction.

We introduce `SimMapNet`, a Bayesian framework that estimates the precision matrix, which serves as the adjacency matrix in a Gaussian graphical model (GGM) for GRN inference. In this approach, the precision matrix—i.e., the inverse of the covariance matrix— represents the network structure, where non-zero entries indicate regulatory relationships between genes [15]. To estimate the precision matrix, we adopt a Bayesian approach, incorporating GO similarities into the estimation of the hyperparameters of the prior distribution. Specifically, GO similarities help to define the prior covariance structure, which guides the Bayesian inference process. Previously, we developed F-MAP [16], which combined Gaussian graphical models with a Bayesian framework to infer the GRN of one species while incorporating gene expression data from related species. In F-MAP, external information was integrated by estimating hyperparameters in the prior distribution through factor analysis of the covariance matrix of related species. Following a similar conceptual framework, `SimMapNet` introduces a distinct methodology by incorporating GO-based functional relationships through kernel functions. Unlike F-MAP, which relies on cross-species covariance structures, `SimMapNet` directly integrates functional similarity measures into the prior distribution, enabling GO similarities to systematically refine the inferred network structure. We evaluate `SimMapNet` using three different microarray gene expression datasets. Two datasets belong to the SOS DNA-repair response pathway in *Escherichia coli*, one with 9 samples and the other with 466 samples, representing different sample sizes of a small-scale GRN [17, 18, 19]. The third dataset, from *Drosophila melanogaster* [20], is considered due to its high dimensionality, which allows us to assess `SimMapNet`’s performance on large-scale data. The results demonstrate `SimMapNet`’s ability to more accurately identify functionally related genes. This novel approach results in a more robust and biologically interpretable network by incorporating functional relationships. Compared to established methods, including Graphical Lasso `(GLASSO)` [21], `KBOOST` [22], and `GENIE3` [23], `SimMapNet` exhibits superior performance in ensuring biologically relevant connections among genes.

## Methods

`SimMapNet` constructs the GRN within the Gaussian Graphical Models (GGM) framework [15], assuming gene relationships follow a multivariate normal distribution, and estimates the precision matrix. The algorithm integrates Bayesian inference and kernel methods to estimate the precision matrix, enforce sparsity and then build adjacency matrices representing regulatory relationships. The Bayesian estimation, GO similarities, and the stepwise implementation of `SimMapNet` are detailed in the following subsections, with additional information on the Bayesian framework and GGM in the supplementary file.

### Bayesian Inference of Precision Matrix

*Let Y* = (*Y*_1_, …, *Y*_*n*_)^*T*^ be an *n × p* matrix, where each *Y*_*i*_, for *i ∈ {*1, …, *n}*, is an independent and identically distributed multivariate normal observation, *Y*_*i*_ *∼ 𝒩* (0, Θ^*−*1^). Here, Θ^*−*1^ is an unknown *p×p* covariance matrix, and Θ is the corresponding precision matrix, which is assumed to be positive definite.

#### Wishart Prior Distribution

The Inverse Wishart distribution is a commonly used prior for Θ^*−*1^ [24, 25]. Based on the relationship between the Wishart and inverse Wishart distributions, the prior distribution on Θ is the Wishart distribution: *W* (*ν*, (*ν*Ω)^*−*1^), where *ν* is the degrees of freedom, which must be greater than (*p −* 1), and Ω is a *p × p* positive definite matrix [25]. The Wishart distribution is the conjugate prior for the population precision matrix of a multivariate normal distribution. Therefore, the posterior distribution of Θ follows the Wishart distribution, *W* (*ν*^*′*^, (*ν*^*′*^Ω^*′*^)^*−*1^), where:

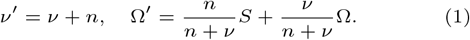

The mode of the posterior distribution, known as the maximum a posteriori (MAP) estimate, serves as an estimator for Θ:

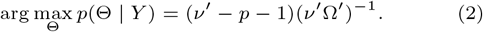

#### Hyperparameter Estimation

The parameter *ν* is emperically determined. It could be learned from the dataset, starting with an initial value greater than *p −* 1, where *p* is the number of variables (genes) [25, 26]. We set the parameter *ν* equals 2*p* recommended by the study of Zhang et al. [27]. To estimate the hyperparameter Ω, GO similarities are transformed into a covariance structure using kernel functions, incorporating gene relationship data. GO similarities range from 0 to 1, and we calculate distances between genes as *d*(*x, x*^*′*^) = 1 *−* similarity, which are then input into the kernel function. Finally, the hyperparameter Ω is estimated as:

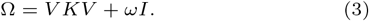

where, *V* = Diag(*σ*_1_, *σ*_2_, …, *σ*_*p*_) is a *p × p* diagonal matrix, whose diagonal entries represent the standard deviations of the corresponding genes. The *p × p* matrix *K* encodes prior information about gene correlations obtained from the kernel function. The matrix *ωI* is a diagonal matrix, where *ω* is a positive parameter that ensures the estimated Ω is positive definite and possesses desirable algebraic properties.

In this study, we focus on two stationary isotropic kernels, which depend only on the distance *d*(*x, x*^*′*^) between inputs [28, 29]. *Squared exponential (SE) kernel* [9, 28, 30] (also known as Gaussian or RBF) and *Ornstein-Uhlenbeck (OU) kernel*, part of the Matérn group [30]. The formulas and parameter definitions for both kernels are provided in the supplementary file (Section 2, subsection “Kernel functions”)..

### Gene Ontology (GO) Similarities

GO classifies gene functions into Biological Process (BP), Molecular Function (MF), and Cellular Component (CC) [12, 31]. GO similarities are computed using semantic similarity measures such as Resnik’s [32] and Wang’s similarity [33](See the supplementary file, Section 3, for more details.). These similarities are mapped to kernel functions to define distances between genes.

#### SimMapNet steps

The steps of the algorithm are presented below. The algorithm is implemented as an R package, available at GitHub. Figure 1 displays a graphical abstract of the `SimMapNet` algorithm.

**Fig. 1.**
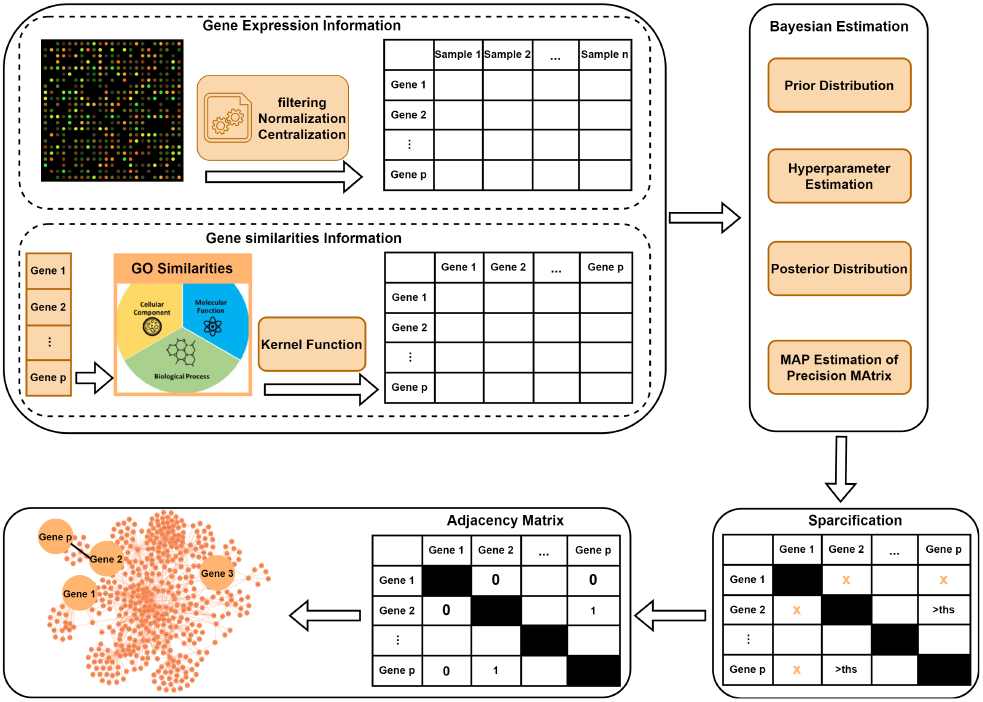
Graphical abstract of the `SimMapNet` algorithm. The “ths” in the Sparsification section denotes the threshold applied to induce sparsity in the MAP estimation of the precision matrix.

##### Step 1: Data Pre-processing

The input to the algorithm consists of gene expression data which is an *n × p* gene expression matrix, where *n* and *p* represent the number of samples and the number of genes, respectively. This matrix can be obtained from various transcriptomic platforms. To ensure the data conforms to a multivariate normal distribution, a normalization transformation is applied. One common transformation is log-transformation, where the log_2_ of the data is computed [34]. The gene expression vectors for each sample are then centered so that each gene has zero mean across samples.

##### Step 2: Similarities calculation and Kernel computation

Calculate the similarities (or differences) between variables. We calculate the GO similarities between genes using the GOSemSim R package [31]. These similarities are then converted into distances and passed to a kernel function to construct the kernel matrix (*K*).

##### Step 3: Bayesian inference of the precision matrix

This step includes multiple stages:

3.1 Set a Wishart prior for the precision matrix: *W* (*ν*, (*ν*Ω)^*−*1^).

3.2 Choose a value equal to (or greater than) 2*p* as the prior degree of freedom *ν*.

3.3 Estimate the hyperparameter Ω using Equation (3).

3.4 Estimate the posterior distribution: *W* (*ν*^*′*^, (*ν*^*′*^Ω^*′*^)^*−*1^) using Equation (1).

3.5 Estimate the MAP of the precision matrix using Equation (2).

##### Step 4: Make Sparse Precision Matrix

To induce sparsity in the MAP estimate, we apply a hard-thresholding method. In this process, different percentiles of the absolute values of the estimated partial correlations are used as thresholds.

##### Step 5: Make Adjacency Matrix

The final binary adjacency matrix is derived from the sparse precision matrix, where non-zero elements (set to one) indicate the presence of an edge between the corresponding gene pairs.

## Results

The results of implementing `SimMapNet` on different datasets are evaluated by following performance metrics, such as True Positive rate (TPR), False Positive rate (FPR), precision (PPV), accuracy (ACC), and F1-score [35]. The mathematical definition of these metrics are provided in supplementary file Section 4. TPR and FPR are also used to plot the receiver operating characteristic (ROC) curve, and the area under the ROC curve (AUC) is calculated. Similarly, the Precision-Recall (PR) curve is plotted using PPV and TPR, and the area under this curve (PRAUC) is calculated [36, 37]. The performance of `SimMapNet` is benchmarked against three well-known methods: GENIE3 [23], KBOOST [22], and GLASSO [21]. In addition, to more precisely evaluate the role of GO similarities, Euclidean distances have been used as prior information for calculating the kernel functions, following the approach described in the `SimMapNet` methodology. The parameters of the algorithm and other methods are trained based on their corresponding diagnostic measures, specifically the F1-score. The optimal parameter set is selected by maximizing the F1-score. To ensure the reliability and robustness of `SimMapNet`, we systematically evaluated its performance using bootstrap datasets and computed the 95% confidence interval (CI) for the F1-score obtained through bootstrap sampling [38]. The CIs are calculated according to the percentile method using percentiles of the bootstrap distribution; 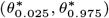. These evaluations allowed us to assess the stability of `SimMapNet` in reconstructing GRNs and its sensitivity to variations in input data.

### SOS DNA repair network dataset

The SOS DNA repair network is a small-scale, experimentally validated gene expression dataset consisting of nine genes. There are two real microarray SOS datasets, varying in terms of their sample size (referred here as SOS1 and SOS2). SOS1 contains nine samples [17, 18], while SOS2 (version 4, build 6) contains 466 samples for the same nine genes [19]. The reference network includes 24 regulatory interactions (edges). GO similarities for MF and BP were mapped for eight genes. As a result, the edges associated with *umuDC* were removed from the reference network, yielding a final network with 22 edges. Figure 2 displays the SOS reference networks, labeled “Reference Network”, along with the reconstructed networks using GO similarities MF (MF-GO), GO similarities BP (BP-GO) for SOS1 and SOS2. Due to the absence of CC similarity data for some genes, the network construction based on CC similarity was performed separately. In the reference network, edges lacking CC information were excluded, resulting in the reference network (`SimMapNetCC`), which contains only eight edges (see Figure S1 for the Reference Network). The constructed GRN using CC similarities are in Figure S1. In addition, the performance metrics for both datasets are in Table 1 and Table 2. Figure 3 displays box plots of the F1-scores achieved from 100 bootstrap sampling for different methods. The results show that `SimMapNet`, particularly with molecular function (`SimMapNet MF`) and biological process (`SimMapNet BP`) GO similarities, achieves the highest F1-scores across both SOS1 and SOS2 datasets, outperforming GLASSO, GENIE3, and KBOOST. The boxplots of bootstrap sampling for both SOS1 and SOS2 confirm `SimMapNet` robustness, as it maintains a high median F1-score with reasonable variability.

**Fig. 2.**
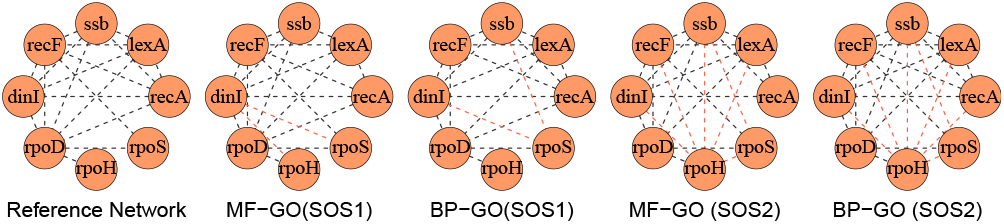
SOS Gene regulatory networks based on MF and BP similarities. The Reference Network represents the true network where each node shows a gene and each edge shows the true relationship between genes. True positive edges are in black and false positives are in red. MF-GO (SOS1) and BP-GO (SOS1) are the reconstructed networks for SOS1 using `SimMapNet` with MF and BP GO similarities, respectively. Similarly, MF-GO (SOS2) and BP-GO (SOS2) are for SOS2.

**Table 1.**
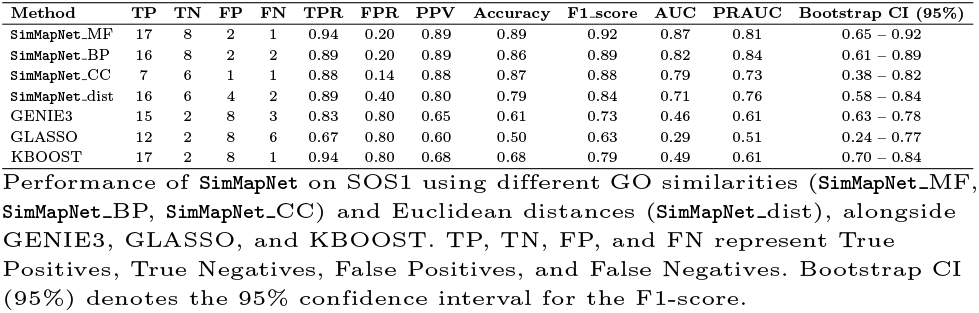
Performance Metrics of Different Methods for SOS1 Constructed Networks

**Table 2.** Performance Metrics of Different Methods for SOS2 Constructed Networks

| Method | TP | TN | FP | FN | TPR | FPR | PPV | Accuracy | F1_score | AUC | PRAUC | Bootstrap CI (95%) |
| --- | --- | --- | --- | --- | --- | --- | --- | --- | --- | --- | --- | --- |
| SimMapNet_MF | 16 | 4 | 6 | 2 | 0.89 | 0.60 | 0.73 | 0.71 | 0.80 | 0.66 | 0.71 | 0.70 - 0.80 |
| SimMapNet_BP | 17 | 3 | 7 | 1 | 0.94 | 0.70 | 0.71 | 0.71 | 0.81 | 0.60 | 0.70 | 0.71 - 0.81 |
| SimMapNet_CC | 7 | 3 | 4 | 1 | 0.88 | 0.57 | 0.64 | 0.67 | 0.74 | 0.66 | 0.60 | 0.53 - 0.84 |
| SimMapNet_dist | 17 | 2 | 8 | 1 | 0.94 | 0.80 | 0.68 | 0.68 | 0.79 | 0.66 | 0.71 | 0.72 - 0.84 |
| GENIE3 | 11 | 7 | 3 | 7 | 0.61 | 0.30 | 0.79 | 0.64 | 0.69 | 0.58 | 0.75 | 0.56 - 0.69 |
| GLASSO | 13 | 8 | 2 | 5 | 0.72 | 0.20 | 0.87 | 0.75 | 0.79 | 0.71 | 0.83 | 0.66 - 0.79 |
| KBOOST | 16 | 3 | 7 | 2 | 0.89 | 0.70 | 0.70 | 0.68 | 0.78 | 0.70 | 0.81 | 0.68 - 0.78 |

Performance of SimMapNet on SOS2 using different GO similarities (SimMapNet\_MF, SimMapNet\_BP, SimMapNet\_CC) and Euclidean distances (SimMapNet\_dist), alongside GENIE3, GLASSO, and KBOOST. TP, TN, FP, and FN represent True Positives, True Negatives, False Positives, and False Negatives. Bootstrap CI (95%) denotes the 95% confidence interval for the F1-score.

**Fig. 3.**
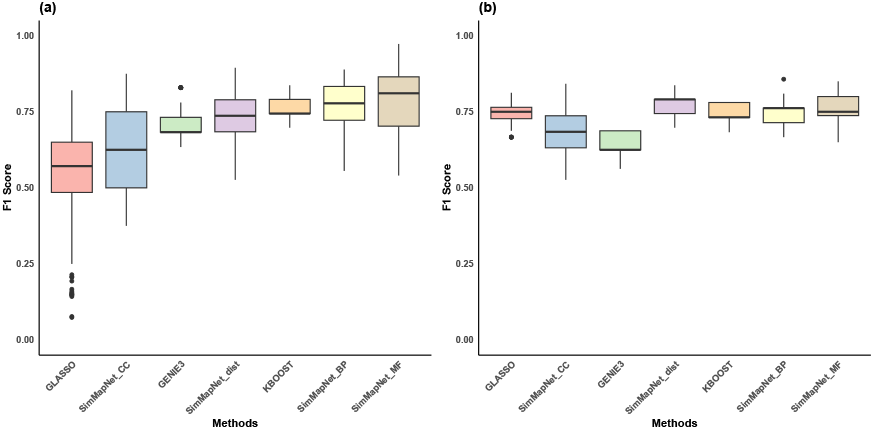
The box plots show the F1-scores obtained from 100 bootstrap samples for different methods, where (a) corresponds to SOS1 and (b) to SOS2. `SimMapNet` is evaluated using GO similarities in Molecular Function (MF), Biological Process (BP), and Cellular Component (CC), denoted as `SimMapNet MF, SimMapNet BP`, and `SimMapNet CC`, respectively. `SimMapNet` dist refers to the version of `SimMapNet` that incorporates Euclidean distances between gene expressions. For comparison, the results of three other methods—GLASSO, KBOOST, and GENIE3—are also included.

Comparing the results of SOS1 and SOS2 reveals that the number of false positive edges in SOS2 is higher than in SOS1, possibly due to greater expression variability resulting from its larger sample size. To assess the impact of sample size on the performance of the constructed networks, we conducted a benchmark on SOS2 using different sample sizes (20, 50, 100, and 200). The samples were selected based on their variability in gene expression. Specifically, we sorted the samples according to their standard deviation across all genes and selected the top *n* samples with the highest standard deviations. Figure 4 displays the changes in F1-score and AUC for different sample sizes across various methods. The complete results are in Table S1.

**Fig. 4.**
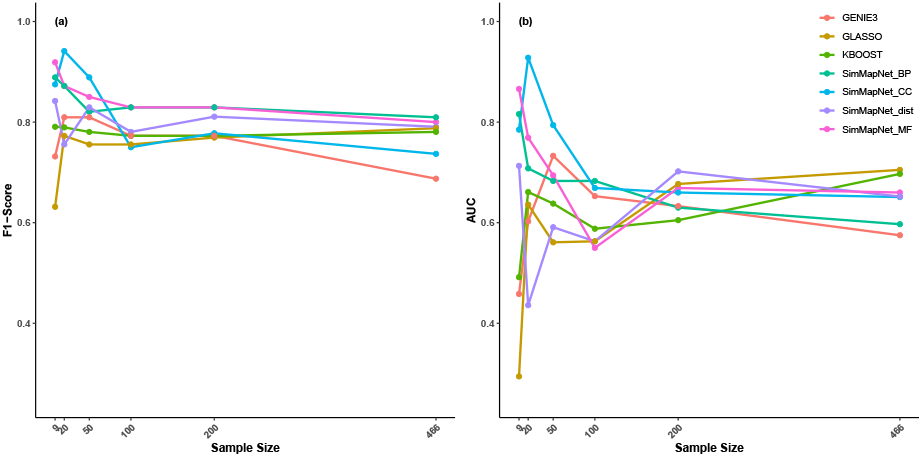
(a) The trend of changes in F1-score of the SOS network across different sample sizes for various methods. (b) The trend of changes in AUC of the SOS network across different sample sizes for various methods. `SimMapNet` using GO similarities MF, BP, and CC are represented as `SimMapNet MF, SimMapNet BP, and SimMapNet CC`, respectively. `SimMapNet dist` represents the `SimMapNet` method using Euclidean distances between gene expressions. Other methods are indicated by their names: GLASSO, KBOOST, and GENIE3. A sample size of 9 corresponds to the results from SOS1 data, while the other sample sizes reflect results from sample selections in SOS2. The sample size of 466 represents the results of the main SOS2 dataset (whole information).

### Drosophila melanogaster Dataset

To assess `SimMapNet`’s performance on high-dimensional datasets, we applied it to gene expression data from *D. melanogaster* (amel), obtained from Kalinka et al. [20] (ArrayExpress, E-MTAB-404). The dataset includes over 3,000 genes, but we focus on those in the reference network, which involves 12 transcription factors (TFs) and their target genes. The reference network is derived from ChIP-chip data from MacArthur et al. [39], which includes relationships between 12 TFs and their 2,049 target genes. In other hand, since GO similarities are unavailable for all genes, we limited the study to 1,441 genes with available all GO similarity information. Genes without GO similarity were excluded from both the study and the reference network.

The best performances for the *Drosophila melanogaster* datasets are shown in Table S2. Networks constructed using `SimMapNet` with GO similarities consistently achieved higher F1-scores than those built by other methods. Each network’s parameters were selected based on the configuration yielding the maximum F1-score, resulting in networks with varying levels of sparsity. To ensure a fair comparison, we searched each method’s results to find the performance corresponding to networks with a similar number of edges to the reference network (about 4,967 edges). Table 3 presents these results. Even under matched conditions, `SimMapNet` using GO similarities clearly outperforms the competing methods.

**Table 3.** Performance Metrics of Different methods for *Drosophila melanogaster* Constructed Networks with similar number of edges

| Method | TP | TN | FP | FN | TPR | FPR | Precision | Accuracy | F1 Score |
| --- | --- | --- | --- | --- | --- | --- | --- | --- | --- |
| SimMapNet_MF | 1992 | 9290 | 2973 | 2959 | 0.40 | 0.24 | 0.40 | 0.66 | 0.40 |
| SimMapNet_BP | 1532 | 8824 | 3439 | 3419 | 0.31 | 0.28 | 0.31 | 0.60 | 0.31 |
| SimMapNet_CC | 2076 | 9366 | 2897 | 2875 | 0.42 | 0.24 | 0.42 | 0.66 | 0.42 |
| SimMapNet_dist | 1436 | 8732 | 3531 | 3515 | 0.29 | 0.29 | 0.29 | 0.59 | 0.29 |
| GENIE3 | 1431 | 8798 | 3465 | 3520 | 0.29 | 0.28 | 0.29 | 0.59 | 0.29 |
| KBOOST | 1460 | 8717 | 3546 | 3491 | 0.29 | 0.29 | 0.29 | 0.59 | 0.29 |
| GLASSO | 82 | 12074 | 189 | 4869 | 0.02 | 0.01 | 0.30 | 0.71 | 0.03 |

Performance comparison of different networks for *Drosophila melanogaster* data, ensuring a similar number of edges across methods. The table includes SimMapNet with different GO similarities (SimMapNet\_MF, SimMapNet\_BP, SimMapNet\_CC) and Euclidean distances (SimMapNet\_dist), as well as GENIE3, GLASSO, and KBOOST. TP, TN, FP, and FN denote True Positives, True Negatives, False Positives, and False Negatives. Bootstrap CI (95%) indicates the confidence interval for the F1-score.

To investigate whether combining networks enhances performance, we integrated the networks constructed using different GO similarity measures by identifying common edges among them (Figure 5, see also Table S3). The bar plot illustrates performance across various GO combinations. The MF&CC network achieved the highest TPR, while MF&BP showed superior Precision, and BP&CC offered a balanced trade-off between the two. Combining all three GO types (MF&BP&CC) resulted in a lower FPR and improvements in both Precision and Accuracy, but with a reduction in TPR. This indicates a trade-off: enhancing precision may come at the expense of detecting fewer true positive edges.

**Fig. 5.**
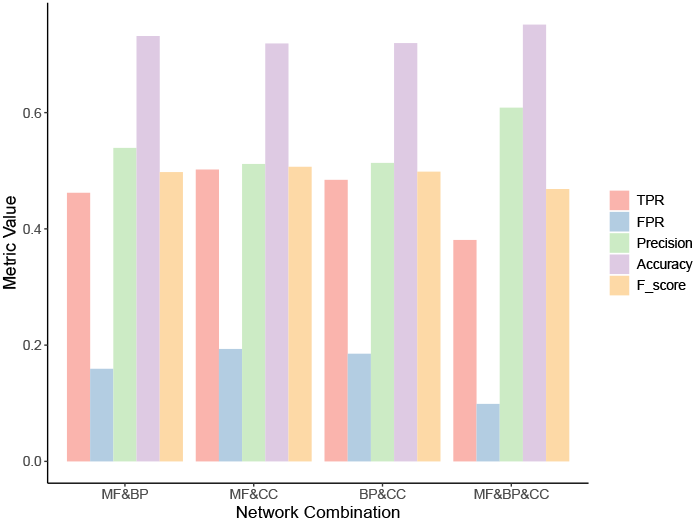
Performance comparison of gene regulatory networks constructed using `SimMapNet` with different GO similarity measures in *Drosophila melanogaster*. The bar plot presents True Positive Rate (TPR), False Positive Rate (FPR), Precision, Accuracy, and F-score for networks based on pairwise similarities (MF&BP, MF&CC, BP&CC) and the combined network (MF&BP&CC). The MF&BP&CC network, which integrates all three GO similarities, achieves the highest accuracy but a lower TPR compared to pairwise networks.

## Discussion

This study introduces `SimMapNet`, a GRN reconstruction approach that uses GO similarity to improve gene relationship inference. Three datasets were used to evaluate `SimMapNet`: two small SOS networks [17, 19] and a large-scale *Drosophila melanogaster* dataset [20]. We found that GO similarities greatly increase network inference accuracy. Further, the results of bootstrap sampling confirm that `SimMapNet` balances both high accuracy and stability, making it a more reliable approach for network inference. The inferred networks were validated against reference networks and benchmarked against GENIE3, KBOOST, and GLASSO. `SimMapNet` typically outperformed various algorithms, with improvements varying by similarity metric. GO-based similarities were more accurate than Euclidean distances, showing the value of biological context over statistics.

Our key assumption is that genes sharing functional similarity are likely to participate in related biological processes and be regulated by similar mechanisms. This often results in potential interactions within a gene regulatory network, where one gene can directly influence the expression of another, enabling coordinated control over a common function [12]. Incorporating such similarities allows the algorithm to better capture the underlying biological structure and mitigate noise in gene expression data, particularly in cases with limited sample sizes. We also incorporate CC similarities, as they may provide complementary insights into gene interactions, despite their implications differing from those of functional similarity. Our results suggest that functional similarity, particularly utilizing MF and BP, generally improves network inference performance. In the construction of SOS GRNs using both SOS1 and SOS2 datasets, networks built with BP and MF similarities outperform those based on CC similarity. In the *Drosophila melanogaster* dataset, the differences are less pronounced, although the constructed network utilizing MF and BP still achieves slightly higher AUC scores than CC (Figure 2 and Figure S1). To ensure a fair comparison, we selected parameter sets that generated a comparable number of network edges, approximately matching the reference network’s edge count (Table 3). Under these conditions, `SimMapNet` using MF remains the strongest predictor, while CC outperforms BP. These findings suggest that while functional similarity is generally more informative for GRN reconstruction, CC similarity may still provide valuable insights depending on the dataset and network structure. Further investigation is needed to fully understand the implications of this result.

Assessment of `SimMapNet`’s performance across varying sample sizes for the SOS dataset (Table S1) reveals distinct performance patterns across GO categories. `SimMapNet` using MF similarities demonstrates the most stable performance, particularly excelling in small sample sizes where functional constraints are more informative. The GRN constructed by `SimMapNet` applying BP performs comparably but shows slight variations due to the complexity of process-level interactions. `SimMapNet` using CC exhibits robust performance but is more sensitive to increasing sample sizes, likely due to structural dependencies captured in cellular localization data. Despite these variations, all GO-constrained versions of `SimMapNet` outperform traditional methods across the tested sample sizes. As the sample sizes increase, we observe that `SimMapNet`, like other GRN inference methods, produces slightly denser networks, which may introduce additional false positives. This effect is common in network reconstruction due to the increasing number of detectable correlations in large datasets. However, `SimMapNet`’s Bayesian framework mitigates this issue better than alternative approaches by integrating biological priors. The results indicate that `SimMapNet` does not require a large sample size to perform well, which makes it particularly effective when data are limited.

GENIE3 performs well at smaller sample sizes but fails to show consistent improvement as sample sizes increase, suggesting that its tree-based structure may be more sensitive to data variability. While GLASSO maintains relatively stable, though suboptimal, performance across larger sample sizes, it is worth mentioning that GLASSO often struggles with overfitting or unstable covariance estimates in low-data scenarios [40, 41]. However, `SimMapNet` outperforms KBOOST not only when applied to SOS data with varying sample sizes but also in constructing gene regulatory networks for Drosophila flies, particularly benefiting from GO-based constraints that improve stability when data is limited.

`SimMapNet` employs a Bayesian framework to estimate the precision matrix of gene expression data, which serves as the foundation for inferring a GGM of the gene regulatory network. While Bayesian precision matrix estimation for multivariate normal data is a well-established and widely used technique [42, 43, 44, 11], most existing approaches are not specifically tailored for GRN construction and lack the capacity to incorporate external biological information. The supplementary file, section 6 and Table S4, includes the results of implementing one of these Bayesian methods—Ledoit and Wolf [42]—on SOS and *Drosophila melanogaster* datasets.) Integrating this information into the prior distribution not only introduces an adaptive regularization mechanism that enhances the interpretability of inferred networks but also reduces computational complexity. As a result, `SimMapNet` achieves remarkably short computation times, making it a highly efficient tool for GRN inference, capable of constructing even large networks within a minute, provided that gene similarities are available. Future studies could explore hybrid approaches that combine Bayesian shrinkage techniques with biological constraints to further enhance GRN inference accuracy.

To incorporate GO similarities, we employ a kernel function that transforms similarity values into a structured prior covariance matrix. This kernel-based approach ensures that prior biological knowledge is treated as a probabilistic influence rather than a strict constraint, making the framework more flexible and biologically realistic. We utilize two isotropic kernel functions: the squared exponential (SE) kernel, which allows smooth transitions in similarity contributions and demonstrated robust performance [45, 9], and the Ornstein-Uhlenbeck (OU) kernel, which enforces a stronger locality constraint to reduce false positive edges [28]. The choice of the kernel function is left to the user, who can select the one that yields the best-performing network. In this study, we evaluate both kernels and determine which one provides the most biologically meaningful results.

The other parameters of `SimMapNet`, such as the degree of freedom of the prior distribution (*ν*), must be specified by the user. Here, we set *ν* = 2*p* since varying it did not strongly affect the results. In this paper, our primary goal is to present `SimMapNet` to introduce an approach that can benefit from incorporating external biological information. Therefore, we optimize the parameters to achieve the best performance, particularly focusing on the F1-score using the available reference network. We select the level of sparsity that maximizes network accuracy rather than using predefined statistical criteria. However, users can apply alternative sparsity selection methods, such as the Bayesian Information Criterion (BIC) [46] or Extended BIC (EBIC) [47, 43], depending on their specific dataset and application. Additionally, we are working on deriving a closed-form solution for certain parameters, particularly leveraging the algebraic properties of the estimations. This effort aims to reduce user dependency on manual parameter selection, thereby improving the usability and robustness of the framework.

While this study focuses on GRN inference, the underlying framework of `SimMapNet` is highly adaptable to other biological datasets, such as microbiome networks. In addition, by leveraging different types of domain-specific similarities, `SimMapNet` has the potential to improve network inference across diverse biological systems.

## Author contributions statement

MSH and MS developed the idea. MSH collected the data. MSH and RA led all statistical analyses from data preprocessing to fitting models, as well as summarizing the results by the creation of figures. MSH wrote the initial complete draft of the manuscript. MSH and RA and MS contributed in interpretations, editing, and revision of the manuscript. All authors read and approved the final manuscript.

## Acknowledgments

This research was supported by the Iran National Science Foundation (INSF) under project number 4027812.

