## Supplementary file for "SimMapNet: A Bayesian Framework for Gene Regulatory Network Inference Using Gene Ontology Similarities as External Hint"

Maryam Shahdoost[1], Roza Aghdam[1], and Mehdi Sadeghi[2]

[1]School of Biological Sciences, Institute For Research In Fundamental Sciences(IPM), Tehran, Iran
[2]Department of Medical Genetics,National Institute for Genetic Engineering and Biotechnology,Tehran,Iran

#### 1 Gaussian Graphical models(GGM)

Graphical models are statistical frameworks where a graph visually represents the conditional dependence structure between variables [1]. Among these, GGMs are widely used to model conditional dependence relationships among variables by leveraging their joint distribution, assuming the variables follow a multivariate normal distribution [2]. In GGMs, the precision matrix ($\Theta$) serves as a key representation. Specifically, each element of the precision matrix ($\theta_{ij}$) quantifies the partial correlation ($\rho_{ij}$) between two corresponding genes, calculated as:

$$\rho_{ij} = \frac{\theta_{ij}}{\sqrt{\theta_{ii}\theta_{jj}}} \tag{1}$$

This implies that genes $i$ and $j$ are conditionally dependent if the corresponding element in the precision matrix is non-zero. In essence, non-zero elements of the precision matrix reveal interactions between two genes.

#### 2 Bayesian Inference of Precision Matrix

Let $Y_i$ for all $i \in \{1, \ldots, n\}$ be independent and identically multivariate normally distributed observations, $Y_i \sim N(0, \Theta^{-1})$, where the $p \times p$ matrix $\Theta^{-1}$ is an unknown covariance matrix. The likelihood function of the data $Y = (Y_1, \ldots, Y_n)^T$ is:

$$L(\Theta \mid Y) = \prod_{i=1}^{n} p(Y_i \mid \Theta) \propto |\Theta|^{n/2} \exp\left(-\frac{n}{2}\mathrm{tr}(S\Theta)\right), \tag{2}$$

where $\Theta$, the precision matrix, is positive definite. Matrix $S$ is the sample covariance matrix, and the maximum likelihood estimator (MLE) of $\Theta^{-1}$.

**Wishart Prior Distribution:** Assuming $Y \sim N(0, \Theta^{-1})$, where $Y$ is an $n \times p$ matrix, the Inverse Wishart distribution is a commonly used prior for $\Theta^{-1}$ [3, 4]. By the relationship between the Wishart and inverse Wishart distributions, the prior distribution on $\Theta$ is the Wishart distribution [5]. The prior distribution of the Wishart $W(\nu, G)$ is:

$$P(\Theta) = \frac{1}{2^{\nu p/2}|G|^{\nu/2}\Gamma_p\left(\frac{\nu}{2}\right)}|\Theta|^{(\nu-p-1)/2} \exp\left(-\frac{1}{2}\mathrm{tr}(G^{-1}\Theta)\right), \tag{3}$$

where the scale matrix $G$ is a $p \times p$ positive definite matrix, $\Gamma_p(.)$ is a multivariate gamma function, and $\nu$ is the degree of freedom, which must be greater than $(p-1)$[5]. The parameter $G$ can be represented as $G = (\nu\Omega)^{-1}$, where $\Omega$ is a $p \times p$ matrix [4]. Thus:

$$\mathbb{E}[\Theta \mid \Omega, \nu] = \Omega^{-1}. \tag{4}$$

The expectation of the covariance matrix is:

$$\mathbb{E}[\Theta^{-1} \mid \Omega, \nu] = \frac{1}{\nu - p - 1}\Omega. \quad (5)$$

The prespecified structural form for $\Omega$ represents structural information about the prior mean of $\Theta$ and $\Theta^{-1}$. The Wishart distribution is the conjugate prior for the population precision matrix of a multivariate normal distribution. Therefore, the posterior distribution of $\Theta$ follows the Wishart distribution, $W(\nu', (\nu'\Omega')^{-1})$, where:

$$\nu' = \nu + n, \quad \Omega' = \frac{n}{n+\nu}S + \frac{\nu}{n+\nu}\Omega. \quad (6)$$

The mode of the posterior distribution (MAP) can be considered as an estimator for $\Theta$:

$$\arg\max_{\Theta} p(\Theta \mid Y) = (\nu' - p - 1)(\nu'\Omega')^{-1}. \quad (7)$$

According to $\Omega'$, the prior degree of freedom $\nu$ represents the strength of belief about prior hyperparameters. It can be set empirically as any non-negative real number greater than $p - 1$ [3, 6]. Some literature, however, imposes a stricter condition, requiring it to be greater than $2 \times p$ [7, 4, 8].

**Hyperparameter Estimation:** The prior Wishart distribution is parameterized by the hyperparameters $\Omega$ and $\nu$, which are essential for defining the posterior distribution. The parameter $\nu$ is learned from the dataset, starting with an initial value of $p + 1$, where $p$ is the number of genes [4, 8]. We set the parameter $\nu$ equals $2\times$ $p$ as it is recommended in Zhang et al. study [7]. To estimate the hyperparameter $\Omega$, we incorporate external information about gene relationships based on GO similarities. Kernel functions are used to transform GO similarities into a covariance structure that reflects these relationships. GO similarities range from 0 to 1, and we calculate distances between genes as $d(x, x') = 1 -$ similarity, which are then input into the kernel function. Finally, the hyperparameter $\Omega$ is estimated as:

$$\Omega = VKV + \omega I. \quad (8)$$

where, $V = \text{Diag}(\sigma_1, \sigma_2, \ldots, \sigma_p)$ is a $p \times p$ diagonal matrix whose diagonal elements are the standard deviations of genes. The $p\times p$ matrix $K$ reveals the prior information about the gene correlation abotained from kernel function.The matrix $\omega I$ is a diagonal matrix, where $\omega$ is a positive parameter ensuring that the estimated $\Omega$ possesses desirable algebraic properties, such as positive definiteness.

**Kernel functions:** In general, a kernel is a mathematical function that measures the similarity or distance between two inputs in a structured way, mapping them to $\mathbb{R}$ [9, 10, 11]. There are various types of kernel functions [12]. In this study, we focus on two stationary kernels. A kernel is stationary if its properties remain unchanged under translations of the input space [13]. Both kernels utilized in this work belong to the class of isotropic kernels, meaning they depend only on the distance $d(x, x')$ between inputs [14].

*The squared exponential (SE) kernel* [15, 12, 16], also known as the Gaussian, radial basis function (RBF), or exponentiated quadratic kernel, is defined as:

$$k_{\text{SE}}(x, x') = a^2 \exp\left(-0.5\frac{d^2(x, x')}{\alpha^2}\right) \quad (9)$$

Here, $a^2$ is the signal variance, controlling the overall scale of the function values. $\alpha$ is the length scale, determining the smoothness of the resulting function. A smaller $\alpha$ results in rapid changes, while a larger $\alpha$ produces smoother variations. $d^2(x, x')$ represents the squared distance between inputs, calculated as $d(x, x') = 1 -$ similarity in this context.

The SE kernel is suitable for capturing smooth and continuous changes in the relationships between genes, making it ideal for modeling GO similarities.

*The Ornstein-Uhlenbeck* (OU) kernel, part of the Matérn kernel group, is another popular kernel for modeling relationships between data points [16]. It is defined as:

$$k_{\text{OU}}(x, x') = \exp\left(-\frac{d(x, x')}{\alpha}\right) \quad (10)$$

Here, $d(x, x')$ is the distance between inputs, calculated as $d(x, x') = 1 -$ similarity. Parameter $\alpha$ is the length scale, controlling the rate of decay. A smaller $\alpha$ results in faster decay of the correlation, while a larger $\alpha$ allows the correlation to persist over larger distances. The OU kernel is well-suited for scenarios where abrupt changes in relationships may occur, providing complementary insights to the SE kernel.

#### 3 Gene Ontology Similarities

Gene Ontology (GO) provides a standardized classification of gene functions, categorized into three main domains: Biological Process (BP), Molecular Function (MF), and Cellular Component (CC) [17, 18, 19, 20]. Each gene in a dataset is associated with one or more GO terms, which describe its role in these domains. The GO terms help establish functional relationships between genes, which is essential for understanding their interactions and regulatory relationships in biological processes.

To quantify the relationships between genes based on GO terms, similarity measures are employed. These measures compute the similarity between genes by evaluating the overlap of their associated GO terms. There are several ways to calculate GO similarities, and these typically rely on semantic similarity measures, which assess how similar the GO terms are in terms of their meanings and hierarchies. For instance, one common approach is based on the structure of the GO terms' ontologies: genes sharing common ancestors in the GO hierarchy are considered more similar than those without shared ancestors [21]. To calculate GO similarity between two genes, a variety of methods can be applied, such as: Resnik's semantic similarity [22, 23], which is based on the most informative common ancestor of two GO terms. Wang similarity [24], which considers the topology of the GO graph. This method assigns weights to each GO term based on its semantic contribution to its descendants. The similarity between two GO terms is computed by aggregating these weighted contributions along their paths in the GO hierarchy. This approach effectively captures both the hierarchical structure and the biological relevance of GO terms.

Once the GO similarities are computed, they can be transformed into a distance measure for use in kernel functions, ensuring that higher similarity corresponds to smaller distances.

#### 4 Performance Metrics

The results of `SimMApNet` are evaluated by following performance metrics, such as True Positive rate (TPR), False Positive rate (FPR), precision (PPV), accuracy (ACC), and F1-score [25]. Mathematically, they are defined by:

$$\begin{aligned}
&\text{TPR} = \frac{\text{TP}}{\text{TP}+\text{FN}}, \quad \text{FPR} = \frac{\text{FP}}{\text{FP}+\text{TN}}, \quad \text{PPV} = \frac{\text{TP}}{\text{TP}+\text{FP}}, \\
&\text{ACC} = \frac{\text{TP}+\text{TN}}{\text{TP}+\text{FP}+\text{TN}+\text{FN}}, \quad \text{F1-score} = \frac{2\times\text{TP}}{2\times\text{TP}+\text{FP}+\text{FN}}
\end{aligned} \tag{11}$$

where TP, FP, TN, and FN are the numbers of True Positives, False Positives, True Negatives, and False Negatives, respectively. TPR and FPR are also used to plot the receiver operating characteristic (ROC) curve, and the area under the ROC curve (AUC) is calculated. Similarly, the Precision-Recall (PR) curve is plotted using PPV and TPR, and the area under this curve (PRAUC) is calculated [26, 27].

#### 5 Performance Metrics of different Methods for SOS Data

the performance metrics for both SOS datasets are in the Supplementary file Table **??** and **??**. Figure S1 displays the SOS reference networks (Reference Network) along with the reconstructed networks using GO similarities MF (MF-GO) , BP (BP-GO) and CC (CC-GO) for SOS1 and SOS2.

To assess the impact of sample size on the performance of the constructed networks, we conducted a benchmark on SOS2 using different sample sizes (20, 50, 100, and 200). The samples were selected based on their variability in gene expression. Specifically, we sorted the samples according to their standard deviation across all genes and selected the top $n$ samples with the highest standard deviations. Table S1 presents the performance metrics of various methods across different sample sizes.

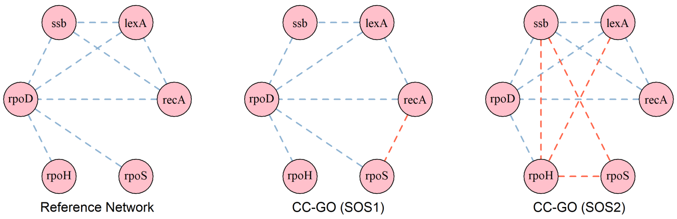


Figure S1: SOS Gene regulatory networks based on CC similarities. The Reference Network represents the true network where each node shows a gene and each edge shows the true relationship between genes. True positive edges are in blue and false positives are in red. SOS1 and SOS2 is representing the reconstructed networks for SOS1 and SOS2.

Table S1: Performance Metrics of Different Methods for SOS data Constructed Networks Across Different Sample Sizes

| Methods | size | TP | TN | FP | FN | TPR | FPR | Precision | Accuracy | F_score | AUC | PRAUC | CI (95%) |
|---|---|---|---|---|---|---|---|---|---|---|---|---|---|
| SimMapNet_MF | 9 | 17 | 8 | 2 | 1 | 0.94 | 0.2 | 0.90 | 0.89 | 0.92 | 0.87 | 0.81 | 0.65–0.92 |
| SimMapNet_MF | 20 | 17 | 6 | 4 | 1 | 0.94 | 0.4 | 0.81 | 0.82 | 0.87 | 0.78 | 0.77 | 0.67–0.87 |
| SimMapNet_MF | 50 | 17 | 7 | 3 | 1 | 0.94 | 0.3 | 0.85 | 0.86 | 0.90 | 0.81 | 0.84 | 0.68–0.95 |
| SimMapNet_MF | 100 | 17 | 5 | 5 | 1 | 0.94 | 0.5 | 0.77 | 0.79 | 0.85 | 0.66 | 0.74 | 0.70–0.85 |
| SimMapNet_MF | 200 | 16 | 4 | 6 | 2 | 0.89 | 0.6 | 0.73 | 0.71 | 0.80 | 0.69 | 0.73 | 0.70–0.85 |
| SimMapNet_MF | 466 | 16 | 4 | 6 | 2 | 0.89 | 0.6 | 0.73 | 0.71 | 0.80 | 0.66 | 0.71 | 0.70–0.80 |
| SimMapNet_BP | 9 | 16 | 8 | 2 | 2 | 0.89 | 0.2 | 0.89 | 0.86 | 0.89 | 0.82 | 0.84 | 0.61–0.89 |
| SimMapNet_BP | 20 | 18 | 7 | 3 | 0 | 1.00 | 0.3 | 0.86 | 0.89 | 0.92 | 0.91 | 0.91 | 0.77–0.92 |
| SimMapNet_BP | 50 | 17 | 4 | 6 | 1 | 0.94 | 0.6 | 0.74 | 0.75 | 0.83 | 0.64 | 0.71 | 0.63–0.83 |
| SimMapNet_BP | 100 | 17 | 4 | 6 | 1 | 0.94 | 0.6 | 0.74 | 0.75 | 0.83 | 0.64 | 0.69 | 0.71–0.83 |
| SimMapNet_BP | 200 | 16 | 4 | 6 | 2 | 0.89 | 0.6 | 0.73 | 0.71 | 0.80 | 0.55 | 0.68 | 0.65–0.80 |
| SimMapNet_BP | 466 | 17 | 3 | 7 | 1 | 0.94 | 0.7 | 0.71 | 0.71 | 0.81 | 0.60 | 0.70 | 0.71–0.81 |
| SimMapNet_CC | 9 | 7 | 6 | 1 | 1 | 0.88 | 0.14 | 0.88 | 0.87 | 0.88 | 0.79 | 0.73 | 0.38–0.82 |
| SimMapNet_CC | 20 | 7 | 6 | 1 | 1 | 0.88 | 0.14 | 0.88 | 0.87 | 0.88 | 0.86 | 0.71 | 0.56–0.88 |
| SimMapNet_CC | 50 | 7 | 4 | 3 | 1 | 0.88 | 0.43 | 0.70 | 0.73 | 0.78 | 0.64 | 0.58 | 0.44–0.77 |
| SimMapNet_CC | 100 | 7 | 5 | 2 | 1 | 0.88 | 0.29 | 0.78 | 0.80 | 0.82 | 0.76 | 0.68 | 0.47–0.82 |
| SimMapNet_CC | 200 | 7 | 4 | 3 | 1 | 0.88 | 0.43 | 0.70 | 0.73 | 0.78 | 0.64 | 0.62 | 0.55–0.78 |
| SimMapNet_CC | 466 | 7 | 3 | 4 | 1 | 0.88 | 0.57 | 0.64 | 0.67 | 0.74 | 0.65 | 0.59 | 0.53–0.84 |
| SimMapNet_dist | 9 | 16 | 6 | 4 | 2 | 0.89 | 0.4 | 0.80 | 0.79 | 0.84 | 0.71 | 0.76 | 0.58–0.84 |
| SimMapNet_dist | 20 | 17 | 6 | 4 | 1 | 0.94 | 0.4 | 0.81 | 0.82 | 0.87 | 0.74 | 0.72 | 0.66–0.87 |
| SimMapNet_dist | 50 | 17 | 2 | 8 | 1 | 0.94 | 0.8 | 0.68 | 0.68 | 0.79 | 0.66 | 0.70 | 0.74–0.84 |
| SimMapNet_dist | 100 | 17 | 4 | 6 | 1 | 0.94 | 0.6 | 0.74 | 0.75 | 0.83 | 0.69 | 0.73 | 0.73–0.83 |
| SimMapNet_dist | 200 | 17 | 4 | 6 | 1 | 0.94 | 0.6 | 0.74 | 0.75 | 0.83 | 0.65 | 0.73 | 0.68–0.83 |
| SimMapNet_dist | 466 | 17 | 2 | 8 | 1 | 0.94 | 0.8 | 0.68 | 0.68 | 0.79 | 0.65 | 0.70 | 0.72–0.84 |
| GLASSO | 9 | 12 | 2 | 8 | 6 | 0.67 | 0.8 | 0.60 | 0.50 | 0.63 | 0.29 | 0.54 | 0.24–0.74 |
| GLASSO | 20 | 17 | 2 | 8 | 0 | 1.00 | 0.8 | 0.68 | 0.72 | 0.80 | 0.50 | 0.58 | 0.34–0.75 |
| GLASSO | 50 | 16 | 3 | 7 | 2 | 0.89 | 0.7 | 0.69 | 0.72 | 0.78 | 0.66 | 0.59 | 0.48–0.76 |
| GLASSO | 100 | 14 | 2 | 8 | 4 | 0.78 | 0.8 | 0.63 | 0.59 | 0.70 | 0.58 | 0.54 | 0.47–0.74 |
| GLASSO | 200 | 16 | 3 | 7 | 1 | 0.94 | 0.7 | 0.69 | 0.71 | 0.77 | 0.65 | 0.58 | 0.50–0.76 |
| GLASSO | 466 | 16 | 5 | 5 | 1 | 0.94 | 0.5 | 0.76 | 0.73 | 0.83 | 0.56 | 0.53 | 0.46–0.71 |
| KBOOST | 9 | 17 | 2 | 8 | 1 | 0.94 | 0.8 | 0.68 | 0.68 | 0.79 | 0.49 | 0.61 | 0.70–0.84 |
| KBOOST | 20 | 15 | 5 | 5 | 3 | 0.83 | 0.5 | 0.75 | 0.71 | 0.79 | 0.66 | 0.70 | 0.58–0.84 |
| KBOOST | 50 | 16 | 3 | 7 | 2 | 0.89 | 0.7 | 0.70 | 0.68 | 0.78 | 0.64 | 0.71 | 0.68–0.83 |
| KBOOST | 100 | 17 | 1 | 9 | 1 | 0.94 | 0.9 | 0.65 | 0.64 | 0.77 | 0.59 | 0.68 | 0.73–0.82 |
| KBOOST | 200 | 17 | 1 | 9 | 1 | 0.94 | 0.9 | 0.65 | 0.64 | 0.77 | 0.60 | 0.69 | 0.73–0.82 |
| KBOOST | 466 | 16 | 3 | 7 | 2 | 0.89 | 0.7 | 0.70 | 0.68 | 0.78 | 0.70 | 0.81 | 0.68–0.78 |
| GENIE3 | 9 | 15 | 2 | 8 | 3 | 0.83 | 0.8 | 0.65 | 0.61 | 0.73 | 0.46 | 0.61 | 0.63–0.78 |
| GENIE3 | 20 | 17 | 3 | 7 | 1 | 0.94 | 0.7 | 0.71 | 0.71 | 0.81 | 0.60 | 0.67 | 0.71–0.86 |
| GENIE3 | 50 | 17 | 3 | 7 | 1 | 0.94 | 0.7 | 0.71 | 0.71 | 0.81 | 0.73 | 0.81 | 0.71–0.81 |
| GENIE3 | 100 | 17 | 1 | 9 | 1 | 0.94 | 0.9 | 0.65 | 0.64 | 0.77 | 0.65 | 0.76 | 0.73–0.82 |
| GENIE3 | 200 | 17 | 1 | 9 | 1 | 0.94 | 0.9 | 0.65 | 0.64 | 0.77 | 0.63 | 0.76 | 0.73–0.82 |
| GENIE3 | 466 | 11 | 7 | 3 | 7 | 0.61 | 0.3 | 0.79 | 0.64 | 0.69 | 0.57 | 0.75 | 0.56–0.69 |

The performance metrics of implementing `SimMapNet` and various methods on the SOS data across different sample sizes. The `SimMapNet` is emplemented using different GO similarities: molecular function (`SimMapNet`_MF), biological process (`SimMapNet`_BP), cellular component (`SimMapNet`_CC), and `SimMapNet`_dist represents the `SimMapNet` method using Euclidean distances between gene expressions. Other methods, including GENIE3, GLASSO, and KBOOST, are also included. The columns TP, TN, FP, and FN refer to True Positive, True Negative, False Positive, and False Negative, respectively. The performance metrics are computed as described in equation 11. Bootstrap CI (95%) represents the 95% confidence interval for the F1-score obtained through 100 bootstrap sampling.

### 6 Performance Metrics of Different Methods for Drosophila fly

The best performances for the Drosophila fly datasets are shown in Table S2. The results show that F1-scores for networks constructed by `SimMapNet` using GO similarities, are higher than those of other methods. The parameter set for each network was chosen based on the highest F1-score, which results in varying network sparsity. However, to provide a fair evaluation, we searched through the results of each method to find the performance corresponding to networks with a similar number of edges to the reference network (approximately 4967 edges). Table **??** presents the results for networks with a comparable number of edges. It is evident that, even under these conditions, `SimMapNet` using GO similarities outperforms the other methods.
To check if the integration of networks can improve the performance, we combined the networks constructed using different Gene Ontology (GO) similarity measures due to find the common edges ( Table S3 ). First, we merged pairs of networks (MF&BP, MF&CC, BP&CC), and then we integrated all three networks (MF&BP&CC). Among the pairwise combinations, the MF&CC network achieved the highest TPR (0.50), while the MF&BP network had the highest Precision (0.54). The BP&CC network showed balanced performance across TPR (0.48) and Precision (0.51). When integrating all three networks (MF&BP&CC), the False Positive Rate (FPR) was significantly reduced (0.1), and Precision improved to 0.61, indicating a more reliable network. However, the TPR decreased to 0.38, suggesting a potential loss of sensitivity. Despite this trade-off, the Accuracy of the final combined network (0.75) was the highest among all models, reflecting an overall improvement in correctly inferred interactions.

Table S2: Performance Metrics of Different Methods for Drosophila fly Constructed Networks

| Method | TP | TN | FP | FN | TPR | FPR | Precision | Accuracy | F1 Score | AUC | PRAUC |
|---|---|---|---|---|---|---|---|---|---|---|---|
| SimMapNet_MF | 3574 | 6425 | 5838 | 1377 | 0.72 | 0.48 | 0.38 | 0.58 | 0.50 | 0.68 | 0.29 |
| SimMapNet_BP | 3094 | 8164 | 4099 | 1857 | 0.62 | 0.33 | 0.43 | 0.65 | 0.51 | 0.69 | 0.29 |
| SimMapNet_CC | 3495 | 6842 | 5421 | 1456 | 0.71 | 0.44 | 0.39 | 0.60 | 0.50 | 0.68 | 0.28 |
| SimMapNet_dist | 4725 | 847 | 11416 | 226 | 0.95 | 0.93 | 0.29 | 0.32 | 0.45 | 0.57 | 0.30 |
| GENIE3 | 4850 | 232 | 12031 | 101 | 0.97 | 0.98 | 0.29 | 0.30 | 0.44 | 0.50 | 0.29 |
| KBOOST | 4925 | 317 | 11946 | 26 | 0.99 | 0.97 | 0.29 | 0.30 | 0.45 | 0.50 | 0.28 |
| GLASSO | 82 | 12123 | 140 | 4869 | 0.02 | 0.01 | 0.37 | 0.71 | 0.03 | 0.0001 | 0.006 |

This table presents the performance of implementing `SimMapNet` for dataset SOS1 using different GO similarities: molecular function (`SimMapNet`_MF), biological process (`SimMapNet`_BP), cellular component (`SimMapNet`_CC), and `SimMapNet`_dist represents the `SimMapNet` method using Euclidean distances between gene expressions. Other methods, including GLASSO, KBOOST and GENIE3 are also included. The performance metrics are computed as described in Equations 11.

Table S3: Performance Metrics of Integrated `SimMapNet` Networks using GO Similarities for Drosophila fly data

| Network | TPR | FPR | Precision | Accuracy | F1 Score |
|---|---|---|---|---|---|
| MF&BP | 0.46 | 0.16 | 0.54 | 0.73 | 0.50 |
| MF&CC | 0.50 | 0.19 | 0.51 | 0.72 | 0.51 |
| BP&CC | 0.48 | 0.19 | 0.51 | 0.72 | 0.50 |
| MF&BP&CC | 0.38 | 0.10 | 0.61 | 0.75 | 0.47 |

This table presents the performance metrics of integrated `SimMapNet` networks using different GO-based similarities. The performance metrics are computed as described in Equations 11.

### 7 Comparing to other Precision Matrix Estimation in Bayesian Framework: Ledoit and Wolf Method

To compare `SimMapNet` with other methods for estimating the precision matrix in Bayesian framework, we implement the shrinkage-based estimator introduced by Ledoit and Wolf [28], referred to here as L&W.
The L&W method is a Bayesian-inspired shrinkage estimator designed to improve the estimation of large covariance matrices, particularly in high-dimensional settings where the number of variables exceeds the number of observations ($p > n$). Instead of adopting a fully Bayesian approach with prior distributions, the method assumes that the true covariance matrix lies between the sample covariance matrix and a structured target matrix (such as an identity or diagonal matrix). The shrinkage intensity, which controls the trade-off between these two components, is optimally computed to minimize estimation error. This approach stabilizes covariance estimation by reducing variance while preserving essential structural

information.
Although the primary application of the L&W method is for high-dimensional data, such as the Drosophila fly dataset analyzed in this study, we also applied it to SOS datasets for comparison. Interestingly, the results for SOS1 and SOS2 are nearly identical, suggesting that when the number of samples is smaller than the number of variables, the L&W method remains relatively insensitive to sample size.
For Drosophila fly data, the `SimMapNet` method outperforms L&W, as shown in Table S4.

Table S4: Performance Metrics of Networks constructed by L&w Method for SOS and Drosophial datasets

| **Method** | **TP** | **TN** | **FP** | **FN** | **TPR** | **FPR** | **Precision** | **Accuracy** | **F1 Score** | AUC | PRAUC |
|---|---|---|---|---|---|---|---|---|---|---|---|
| SOS1 | 17 | 2 | 8 | 1 | 0.94 | 0.8 | 0.68 | 0.68 | 0.79 | 0.66 | 0.71 |
| SOS2 | 17 | 2 | 8 | 1 | 0.94 | 0.8 | 0.68 | 0.68 | 0.79 | 0.68 | 0.79 |
| Drosophila fly | 4925 | 74 | 12189 | 26 | 0.99 | 0.99 | 0.29 | 0.29 | 0.45 | 0.55 | 0.34 |
| Drosophila fly | 1689 | 9063 | 3200 | 3253 | 0.34 | 0.26 | 0.35 | 0.63 | 0.34 | | |

This table presents the performance of implementing Ledoit & Wolf(L&W) method on SOS and Drosophila datasets. For Drosophila fly dataset, there are two rows. The first row represents the best performance network according to F1-Score and the second row indicate the performance metrics for the network including the same number of edges to the reference networks.

### References


[1] Daphane Koller. Probabilistic graphical models: Principles and techniques, 2009.

[2] Ami Wiesel, Yonina C Eldar, and Alfred O Hero III. Covariance estimation in decomposable gaussian graphical models. *IEEE Transactions on Signal Processing*, 58(3):1482–1492, 2009.

[3] Morris L Eaton and ML Eaton. *Multivariate statistics: a vector space approach*, volume 512. Wiley New York, 1983.

[4] Andrew Gelman, John B Carlin, Hal S Stern, and Donald B Rubin. *Bayesian data analysis*. Chapman and Hall/CRC, 1995.

[5] Zhiyong Zhang. A note on wishart and inverse wishart priors for covariance matrix. *Journal of Behavioral Data Science*, 1(2):119–126, 2021.

[6] Chan-Fu Chen. Bayesian inference for a normal dispersion matrix and its application to stochastic multiple regression analysis. *Journal of the Royal Statistical Society: Series B (Methodological)*, 41(2):235–248, 1979.

[7] Yi Zhang. Smart pca. In *Twenty-First International Joint Conference on Artificial Intelligence*, 2009.

[8] S James Press. *Applied multivariate analysis: using Bayesian and frequentist methods of inference*. Courier Corporation, 2005.

[9] Carl Edward Rasmussen and Hannes Nickisch. Gaussian processes for machine learning (gpml) toolbox. *The Journal of Machine Learning Research*, 11:3011–3015, 2010.

[10] Marta Karaliutė and Kestutis Dučinskas. Classification of gaussian spatio-temporal data with stationary separable covariances. *Nonlinear Analysis: Modelling and Control*, 26(2):363–374, 2021.

[11] Sandra De Iaco, Donato Posa, Claudia Cappello, and Sabrina Maggio. On some characteristics of gaussian covariance functions. *International Statistical Review*, 89(1):36–53, 2021.

[12] Andrew Gordon Wilson. *Covariance kernels for fast automatic pattern discovery and extrapolation with Gaussian processes*. PhD thesis, University of Cambridge Cambridge, UK, 2014.

[13] David K Duvenaud, Hannes Nickisch, and Carl Rasmussen. Additive gaussian processes. *Advances in neural information processing systems*, 24, 2011.

[14] Ethan Anderes, Jesper Møller, and Jakob G Rasmussen. Isotropic covariance functions on graphs and their edges. 2020.
